# Evaluation of synthetic data generation for longitudinal 6-year follow-up subclinical cardiovascular disease data using GAN models

**DOI:** 10.64898/2026.10.01.755916

**Authors:** Amanda Bertgren, Fredrik Öhberg, Paolo Soda, Ulf Näslund, Patrik Wennberg, Christer Grönlund

## Abstract

Although largely preventable, cardiovascular disease (CVD) is the leading cause of death worldwide. Synthetic data generation (SDG) of health data can facilitate data sharing for research and development of personalised cardiovascular disease prevention tools. Current state-of-the-art SDG methods are limited to cross-sectional data or focused on short-term or in-patient EHR applications lacking benchmarks for generalisability. Since prevention is a long-term longitudinal objective, these limitations must be addressed.

We evaluated two tabular generative adversarial networks, TGAN and CTGAN, for privacy and fidelity on a structured 6-year longitudinal trial dataset within CVD prevention. We compared performance with a rudimentary perturbation method. In addition, we performed a variable-wise evaluation and an age-stratified analysis of the higher performing model to assess preservation of clinically relevant characteristics.

Overall, CTGAN showed better fidelity than TGAN. The models performed similarly within privacy, showing better results than the perturbation method. CTGAN had lower performance in comparison to the perturbation method within numerical fidelity of continuous variables and overall temporal fidelity. Furthermore, CTGAN’s performance was lower for younger age groups than older.

CTGAN shows better than or equal performance within fidelity and privacy than TGAN and the perturbation method on a long-term longitudinal dataset within CVD prevention. To the best of our knowledge, this is the first study evaluating CTGAN on a longitudinal dataset and our findings suggest that handling of temporal characteristics and imbalanced subgroups need further refinement.

## Introduction

Cardiovascular disease (CVD) is the leading cause of death globally [1]. Lifestyle affects its prevalence, as multiple CVD risk factors are modifiable [2] and even individuals with a hereditary predisposition to CVD can reduce their risk by half with a favourable lifestyle [3]. Therefore, prevention is crucial to reduce the prevalence of CVD. Low perception and underestimation of individual risk is a major barrier to adherence to treatment and lifestyle changes [4], but personalised prevention and intervention can lower this barrier [5, 6].

Personalised prevention can be enhanced by the rise of big data [7] with electronic health records (EHR) in a clinical setting and wearable devices in an ambulatory setting [8]. Digital twins are at the technical forefront of research on personalised cardiovascular healthcare with studies ranging from mechanical heart simulation to tools for monitoring and coaching [9, 10]. Health digital twins can play an important role in prevention, as digital interventions have been shown to increase the adoption of positive habits [11].

The development and research on healthcare digital twins for CVD prevention require access to considerable amounts of data regarding lifestyle, health and surrogate outcomes, raising privacy concerns even after de-identification [12–15]. Compliance with data protection guidelines and regulations such as the General Data Protection Regulation (GDPR) in Europe [16] and the Health Insurance Portability and Accountability Act (HIPAA) in the United States of America [17] impose additional demands on the secondary use of data. People are also reluctant to share their health data collected in an ambulatory setting, even with healthcare professionals, due to concerns about leakage and judgement [18]. Many approaches are used to facilitate the sharing of sensitive data. The removal of identifiers and data manipulation techniques, such as recoding and perturbation, are commonly used in practice [19] despite risking the integrity of the individuals in the dataset [14, 15]. Synthetic data generation (SDG) using generative machine learning models can offer controlled preservation of privacy whilst retaining important characteristics of the cohort and its subgroups [20, 21].

## Related works

Early SDG methods relied on imputation techniques to generate partial tabular datasets [22]. Generative adversarial networks (GAN) was originally introduced to generate synthetic images [23, 24] before being extended to tabular data with MedGAN being among the first GAN models for synthetic health data generation [20].

In general, the adversarial architecture of GANs introduced fundamental challenges during model training due to mode collapse and vanishing gradients [23, 25]. Techniques using the Wasserstein distance metric have shown improved stability during training and contribute to mitigating these challenges for all types of GAN models [26, 27].

In terms of GAN models for tabular data, additional challenges arise. Tabular datasets often include mixed data types, multimodal distributions, and tail-end samples which complicate learning. TGAN and CTGAN address these challenges with techniques that include mode-specific normalisation and conditional sampling methods [28, 29]. CTGAN also incorporates the Wasserstein distance metric and has been repeatedly validated within different types of synthetic EHR generation, as it is commonly used as a benchmark model [21, 30–32], but primarily on cross-sectional datasets, i.e. data that is only measured once. In contrast, health data is conventionally longitudinal, meaning it is measured at multiple points in time.

To address the issue of longitudinal data structure, several models and approaches have been suggested. Synteg simulates ICD codes (codes for International Classification of Diseases) for events that vary in time using a latent space projection of temporal characteristics to address the longitudinal structure [33]. EHR-M-GAN uses a variational autoencoder (VAE) to extract latent space representations on multidomain and longitudinal data from the intensive care unit (ICU) [34]. HALO increases the dimensionality of longitudinal EHR modelling with a hierarchical autoregressive language model for events of inpatient ICU data and outpatient diagnosis code data [35].

Despite these advances in longitudinal SDG research, most state-of-the-art longitudinal approaches remain focused on short-term or disease-prevalent datasets, typically derived from inpatient cohorts. Inpatient data differs from outpatient and non-patient data in structural composition and internal covariance [36]. For non-patient cohorts, such as prevention cohorts, longitudinal records are often sparse, irregularly sampled, which combined with low covariance creates significant analytical challenges [37, 38], potentially resulting in low signal. Since the methods are not widely benchmarked on different types of clinical datasets, their generalisability is not well established, in contrast to their task-specific performance [39]. Therefore, their extendability to longitudinal health data aimed at long-term preventive objectives is uncertain and since longitudinal SDG on health data in general lack standardised evaluation [39], benchmarking is needed for generalisation and comparability.

## Contribution

In this work, we challenge the structural limitations of state-of-the-art cross-sectional methods to contribute to benchmarking within SDG on longitudinal health data, as a first step towards long-term longitudinal SDG in cardiovascular disease prevention. We did this by evaluating the extendability of a cross-sectional SDG method that has been thoroughly benchmarked on health data, CTGAN [29], and its precursor, TGAN [28], on a long-term longitudinal dataset within cardiovascular disease prevention. To the best of our knowledge, this is the first study applying CTGAN and TGAN to a longitudinal dataset. The models were compared to a rudimentary perturbation method.

1. **The primary objective** was to evaluate the models’ performances in fidelity and privacy to identify the highest performing model for generation of synthetic data for longitudinal long-term cardiovascular disease prevention data.
2. **The secondary objective** was to evaluate the preservation of clinically relevant characteristics of the synthetic data for research purposes, by conducting a variable-wise assessment and stratified analysis on age groups of the highest performing model of the primary analysis.

## Materials and methods

The overall study design consists of three main steps: data preprocessing, model training and evaluation.

### Dataset

The dataset was retrieved from the VIPVIZA (VIsualiZAtion of asymptomatic Atherosclerotic disease) study - a randomised controlled trial for individuals at ages 40, 50 and 60 in Västerbotten county, Sweden [5, 40]. The VIPVIZA study investigates the effect of pictorial risk communication on CVD risk as an intervention strategy with participants recruited from Västerbotten Intervention Program (VIP), a screening and prevention programme [41]. Participants undergo an examination that includes questionnaires, measurements, a blood panel and an ultrasound examination of the carotid artery. The examination was performed at baseline and at follow-up after 3 and 6 years and is currently conducting the 10-year follow-up.

Participants were recruited to VIPVIZA based on age-dependent criteria, younger individuals required to meet more stringent risk-related conditions than older participants. Individuals with significant stenosis, as discovered during the ultrasound examination, were excluded from the study and remitted to clinical management, and thus the most extreme cases of pronounced atherosclerosis are not included in the dataset. Hard outcomes on cardiovascular events has not yet been extracted for this cohort but is planned in a couple of years.

The variables in this study spanned multiple health domains from the VIPVIZA study, including clinical risk markers, psychology, ultrasound imaging and lifestyle. The selection of variables was based on the combined recommendation of VIPVIZA experts within the domains. The dataset comprised of time-invariant variables only measured once, such as sex and education level, and time-varying variables measured multiple times, such as blood pressure and body mass index (BMI). A number of *m*_ti_ = 12 time-invariant variables only measured at baseline and *m* = 28 time-varying variables measured at baseline, 3-year and 6-year follow-up (3*m* = 84 variables) were included. We defined the data subsets

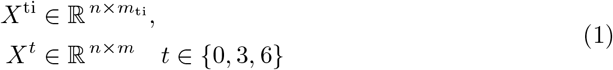

with *X*^ti^ denoting time-invariant variables and *X*^*t*^ each set of time-varying variables. The four subsets were concatenated into one dataset according to

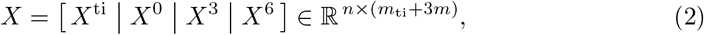

totalling in *m*_ti_ + 3*m* = 96 variables. A graphical overview of the subsets and concatenation is illustrated in Fig 1. The complete dataset (*n*_0_ = 3532) was reduced to *n* = 2612 samples (participants) after removing samples with loss of follow-up to the 3- or 6-year examination. Reasons for loss to follow up are reported in [42, 43]. For completeness, the descriptive statistics of all variables are reported in the supplementary materials S1. The dataset requested for this study was retrieved on March 1, 2024, and all accessed samples were pseudonymised to prevent identification of individuals.

**Fig 1.**
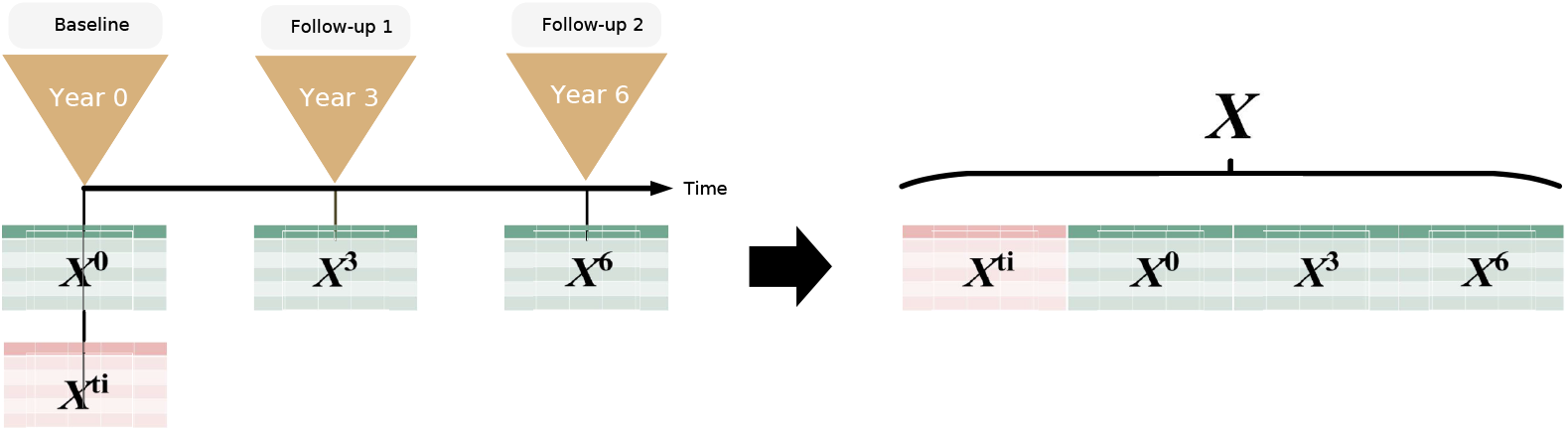
Dataset overview. Graphical overview of time points for collection of data and concatenation of subsets.

The data was split sequentially to avoid information leakage, since the examinations were performed at different points in time. The splits were divided into train (75%), validation (10%) and test (15%). All labels with less than 5% occurrence were merged with other labels and recoded to improve robustness of both imputer and generative models. The effective dimensionality of the dataset after merging and one-hot encoding was 138 variables.

Since participation in the VIPVIZA study is voluntary, any participant could refrain from responding to any part of the questionnaire or partaking in any part of the examination. Despite this, most variables had relatively low levels of missing, except for the variable “*Depression in HADS*” at baseline, which had a notable percentage of missing of 33%. This is because the variable was introduced some time after the start of the study. Since a sequential split is made, this large proportion only affects the training split of the data and may thus have an effect on model performance on the variable.

The missing values were imputed using multiple imputation for mixed data. The imputer was based on miceforest with five mean match candidates and trained on the training data split, then applied on the training, validation and test data. Three imputed datasets were generated to account for stochasticity. A full description of proportion of missing per variable and time point as well as statistics before and after imputation is reported in the supplementary materials S1.

Three datasets, each with a different seed for reproducibility, were generated per model and imputation to account for stochasticity. The results reported were averaged across all imputations and seeded runs.

### Tabular GAN models for synthetic data generation

GANs consist of an adversarial architecture of two directed neural networks [23]. The generator *G* samples random noise *z* distributed over *p*_*z*_(*z*) generating synthetic data 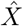 aiming to mimic distribution *p*_*g*_(*X*). The competing discriminator *D* aims to distinguish 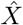 from *X*, that is, minimise *log*(*D*(*x*)) whilst *G* aims to minimise log(1 *™ D*(*G*(*z*))).

TGAN uses conventional cross-entropy loss to optimise *D* and the Kullback-Leibler divergence of numerical and categorical variables is added to *G*s loss function. *G* consists of a long-short-term memory network and *D* of a multilayer perceptron to classify the generated data [28].

CTGAN is based on TGAN. A conditional generator and a train-by-sampling method are included to account for potentially imbalanced categorical variables. Both *G* and *D* consist of fully connected hidden layers due to the lack of local structures inherent in tabular data. *G* consists of residual layers and *D* of linear layers [29].

Both models integrate mode-specific normalisation and one-hot encoding in their respective pipelines [28, 29]. TGAN specifically incorporates a smoothing mechanism for the latter.

### Model training

All hyperparameters were kept at default values except training duration. For fairness, convergence sweeps were performed for both models and identical evaluation procedures were applied. Since CTGAN and TGAN are inherently different in configuration, in which CTGAN employs a conventional GAN batch sampling strategy whilst TGAN employs a strategy in which steps per epoch determines how many samples of batches are drawn per epoch, the search spaces were found by an exploratory sweep per model. The points of convergence were found at 45 000 epochs for CTGAN and 120 epochs with 100 steps per epoch for TGAN.

Both models were trained on an NVIDIA GeForce RTX 2080 Ti GPU.

### Primary evaluation

In our primary evaluation, we evaluated fidelity and privacy, as previously proposed for evaluation of SDG methods [44, 45], to assess the statistical similarity of the datasets and the integrity of the real samples.

#### Fidelity

The fidelity of the synthetic data was evaluated in three areas; univariate, bivariate and temporal fidelity, inspired by previous studies [45, 46].

Univariate fidelity was calculated using the Kolmogorov-Smirnov statistic, *KS*, for numerical variables, and the total variation distance, *TV D*, for categorical variables, to examine distributional distance. Bivariate fidelity was calculated using the difference in Pearson correlation coefficient, *PCD*, for numerical variables, and contingency similarity complement, *CS*^c^, for categorical variables. The metrics were chosen as suggested by [47].

Temporal fidelity was evaluated using the squared maximum mean discrepancy (*MMD*^2^) for the three subsets *X*^0^, *X*^3^ and *X*^6^ of time-varying variables. *MMD*^2^ was selected due to its applicability to variables with few repeated measurements, as is the case in this study with three measurements of the time-varying variables. For studies with a higher number of repeated measurements, metrics such as dynamic time warping and autoregressive likelihood can be considered.

#### Privacy

The privacy of the models was evaluated in two areas; membership and identity disclosure. The Syntheval framework was used for the evaluation of privacy [48].

Membership disclosure refers to the risk that an attacker can infer whether a specific individual was part of the training dataset, evaluated using the epsilon-identifiable risk (*ε*-IR) in this study. Identity disclosure was assessed using nearest neighbour distance ratio (*NNDR*) and median distance to the closest record (*MDCR*) to measure the similarity between real and synthetic samples.

### Secondary evaluation

A secondary evaluation was conducted on the model that achieved the highest performance in the primary assessment to assess preservation of clinically relevant characteristics for the intended use of VIPVIZA synthetic data. The intended use was not limited to one singular downstream task predicting a hard outcome, but rather for the synthetic data to be used in place of original data for research on CVD prevention and health digital twin purposes. Thus, a conventional downstream task evaluation such as train-on-real, test-on-synthetic, would not have effectively assessed whether the clinical relevance of the original data was retained in the synthetic data. Instead, significant clinical properties of the dataset were evaluated in the secondary evaluation. This evaluation consisted of two steps: 1) assessing variable-wise fidelity and 2) performing a stratified analysis of performance metrics across age groups.

#### Variable-wise fidelity

Variable-wise fidelity was assessed across all areas of fidelity. Univariate and temporal fidelity were inherently computed variable-wise, but the bivariate fidelity matrices required aggregation to derive a variable-wise metric. Aggregated bivariate metrics were obtained by computing the L2-norm ( || · ||_2_) of the upper triangular matrices of (*PCD*) and (*CS*^c^) such that

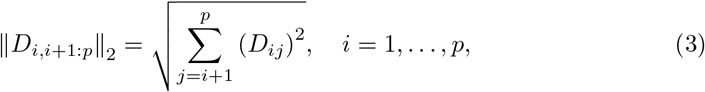

where *D*_*ij*_ is the (*i, j*)-th entry of *D* ∈ {*PCD, CS*^c^}, resulting in an accumulated distance metric.

#### Age-stratified analysis

A stratified analysis across age groups was conducted to evaluate performance for imbalanced subgroups. Age was chosen as the stratifying variable for three reasons; (1) due to well-established health differences between age groups, (2) due to differences in the VIPVIZA inclusion criteria for different ages, thus implicating the presence of three subgroups and (3) due to imbalance in representation of the three subgroups. The analysis was carried out by repeating the computations from the primary analysis for the three age-based strata.

Since the 40-year-olds only constituted 6.9% of the full dataset, we computed its percentage across all splits to verify sufficient occurrence across the splits. Their respective percentages in each of the splits were computed to 7.3% for train, 6.9% for validation and 5.4% for test data.

### Reference perturbation method

Recently, obfuscation and perturbation techniques have been reported to challenge SDG methods in performance, including CTGAN [30, 31]. For this reason, we included a rudimentary method of Gaussian noise addition on the dataset to constitute a point of reference, hereby referenced as the *perturbation method*. The perturbation level was determined based on the trade-off between univariate fidelity and privacy, as follows.

Multiple perturbed datasets were generated by injecting the training dataset with different levels of noise using a parameter *η* ∈ {0.0, 0.1, …, 0.5}. The resulting fidelity and privacy metrics are reported in Fig 2. The types of variables in the dataset were perturbed as follows.

**Fig 2.**
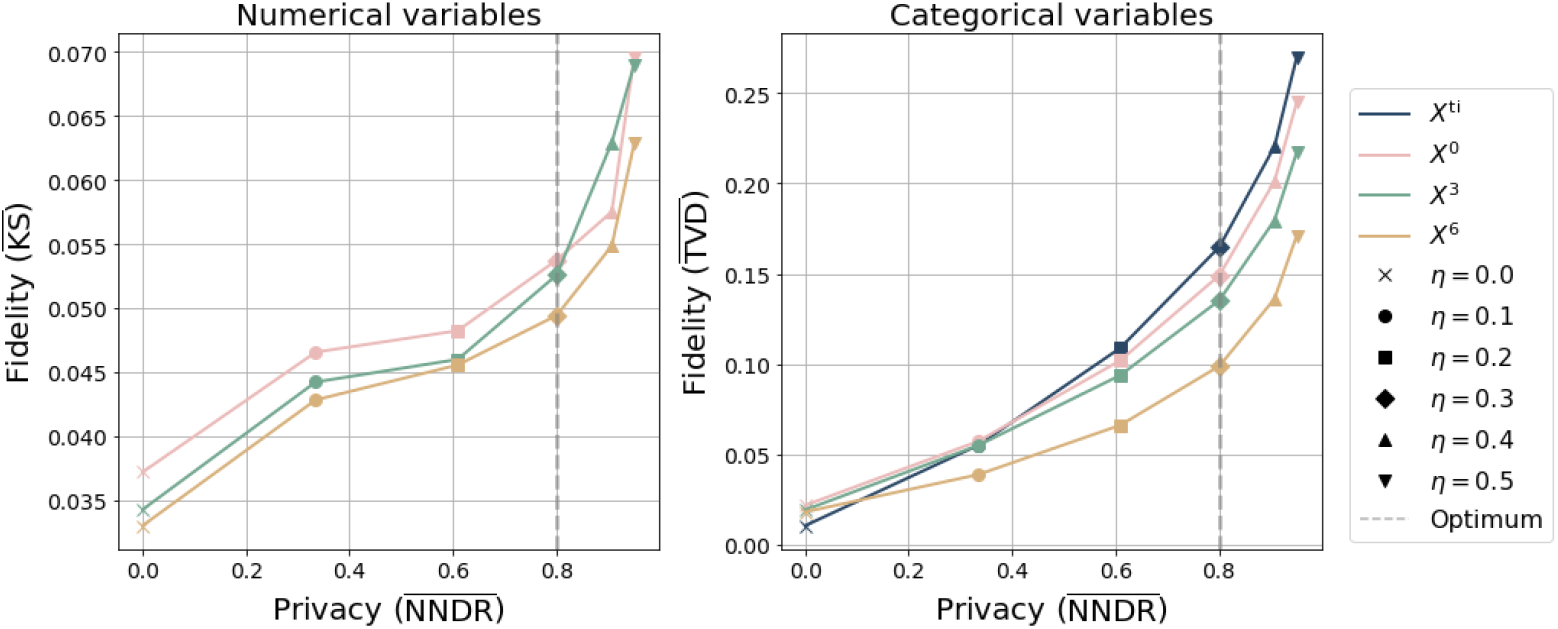
Perturbation method selection. Fidelity versus privacy for perturbed datasets with different levels of noise. The fidelity metrics are mean Kolmogorov-Smirnov statistic 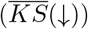 and mean total variation distance 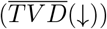 for numerical and categorical variables respectively. The privacy metric is mean distance ratio to nearest neighbour 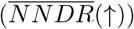. The optimum is computed by taking the mean of the normalised minimum distance to ideal point over all points in time.

- **Numerical variables:** For each value *x*, Gaussian noise was added according to

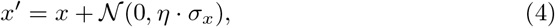

where *σ*_*x*_ denotes the standard deviation of the variable.

- **Categorical variables (excluding booleans):** With probability *η*, the category of a value was replaced by a randomly sampled category from the empirical distribution of the variable.
- **Boolean variables:** With probability *η*, the value was flipped (True ↔ False).

The fidelity-privacy trade-off was evaluated by computing the normalised minimum distance to the ideal point. 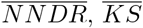, *KS* and 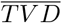 was computed for each level of noise. Note that 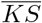 for *X*^ti^ only includes one variable (*Height*) and was thus excluded to avoid skewing the results. Also note that computation of 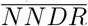for a noise level *η* = 0 results in division by zero since the datasets are identical, and the value was thus clipped to 0.

With (1, 0) being the ideal point after normalisation, the Euclidean distance was computed

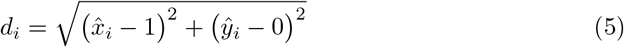

with normalised privacy 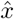 and normalised fidelity *ŷ* according to

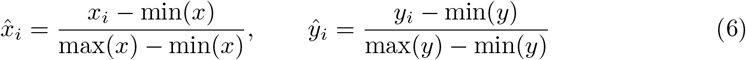

where *x* is 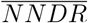 and *y* is 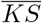 and 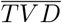, averaged across the time points. The optimums were selected such that

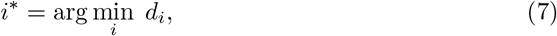

which corresponded to a noise level of *η* = 0.3 for both numerical and categorical variables. This level was therefore adopted for the dataset in the perturbation method. The optimums are reported in Fig 2.

## Results

### Primary evaluation

The fidelity for TGAN, CTGAN and the baseline is shown in Fig 3. Overall, CTGAN and the perturbation method displayed highest performance in different areas of fidelity. CTGAN displayed most consistent performance across all areas. For univariate fidelity, the perturbation method showed highest performance for numerical variables, but CTGAN showed highest performance for categorical variables. The perturbation method showed high variability in performance across numerical and categorical variables. The same trend held for bivariate fidelity, as the perturbation method had highest performance for numerical variables, and CTGAN for categorical variables. The perturbation method showed highest performance on temporal fidelity, followed by CTGAN. There was no general trend across the time-point subsets.

**Fig 3.**
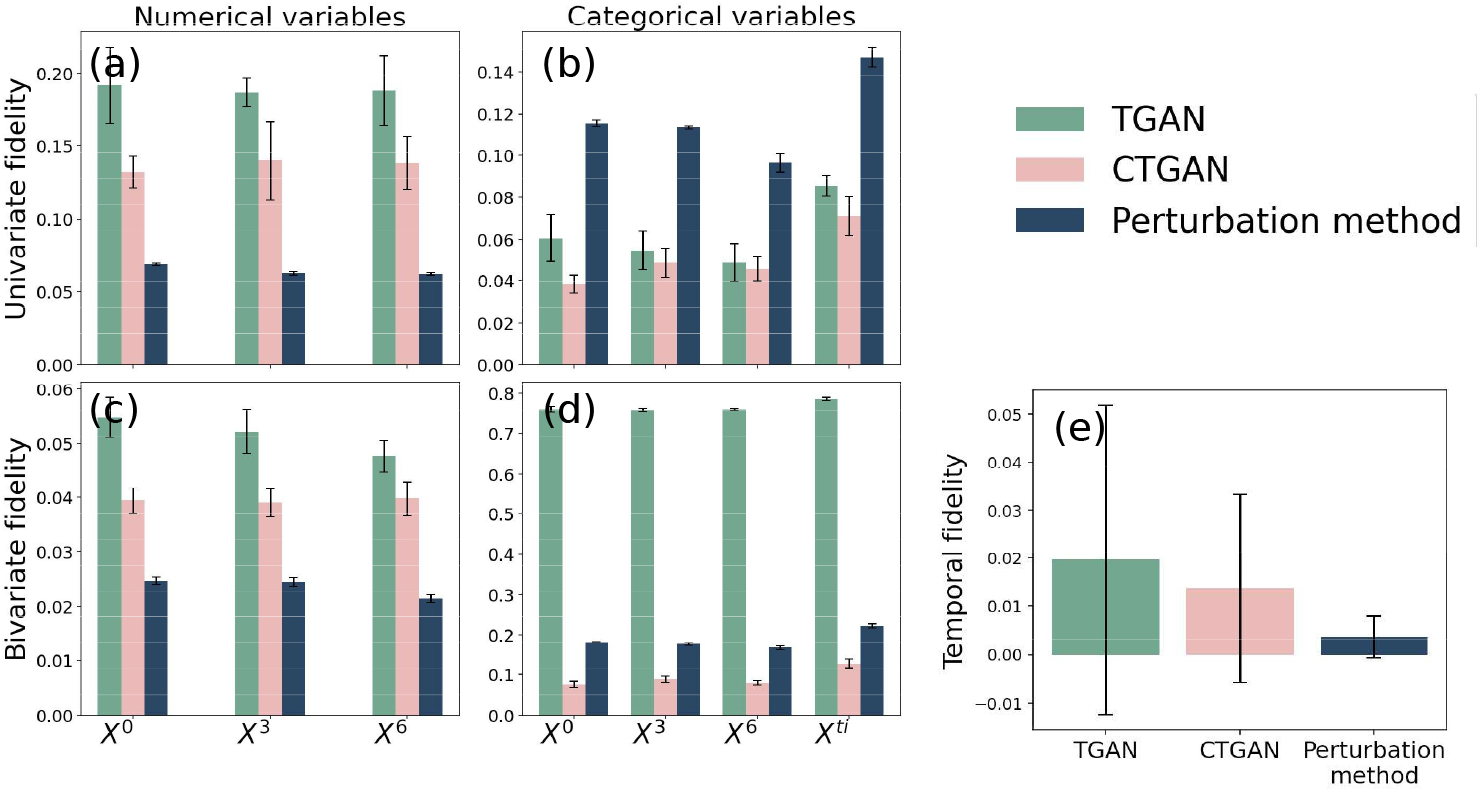
Fidelity. Fidelity for TGAN, CTGAN and baseline. The univariate fidelity includes mean Kolmogorov-Smirnov 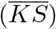 in (a) and mean total variation distance 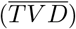 in (b) for numerical and categorical variables respectively. The bivariate fidelity includes mean L2-norm of difference in Pearson correlation coefficient 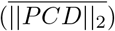 in (c) and mean L2-norm of contingency similarity complement 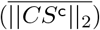 in (d) for numerical and categorical variables respectively. Temporal fidelity includes mean maximum mean discrepancy squared 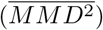 across all points in time, reported in (e). Lower values indicate better performance of all metrics.

The privacy results are reported in Table 1. The performance on identity disclosure metrics (*NNDR* and *MDCR*) were similar for both models but lower for the perturbation method. The high risk of membership disclosure (*ε ™ IR* = 91.3%), using the perturbation method was notable.

**Table 1.**
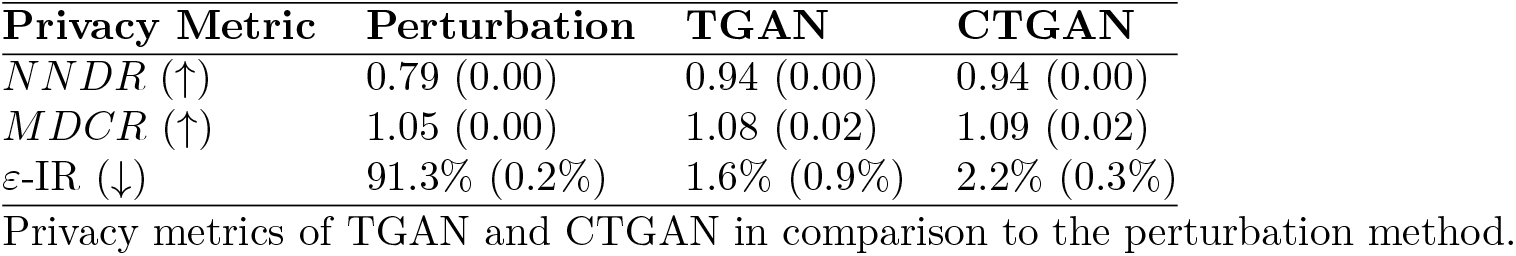
Model privacy.

### Secondary evaluation

CTGAN was selected for the secondary evaluation since it showed better fidelity than TGAN and similar privacy performance to TGAN.

### Variable-wise fidelity

Variable-wise fidelity metrics of CTGAN in comparison to the perturbation method are displayed in Fig 4. Overall, the variable-wise results show similar trends for numerical and categorical variables as displayed in fig. 3 as CTGAN generally performed better for categorical variables.

**Fig 4.**
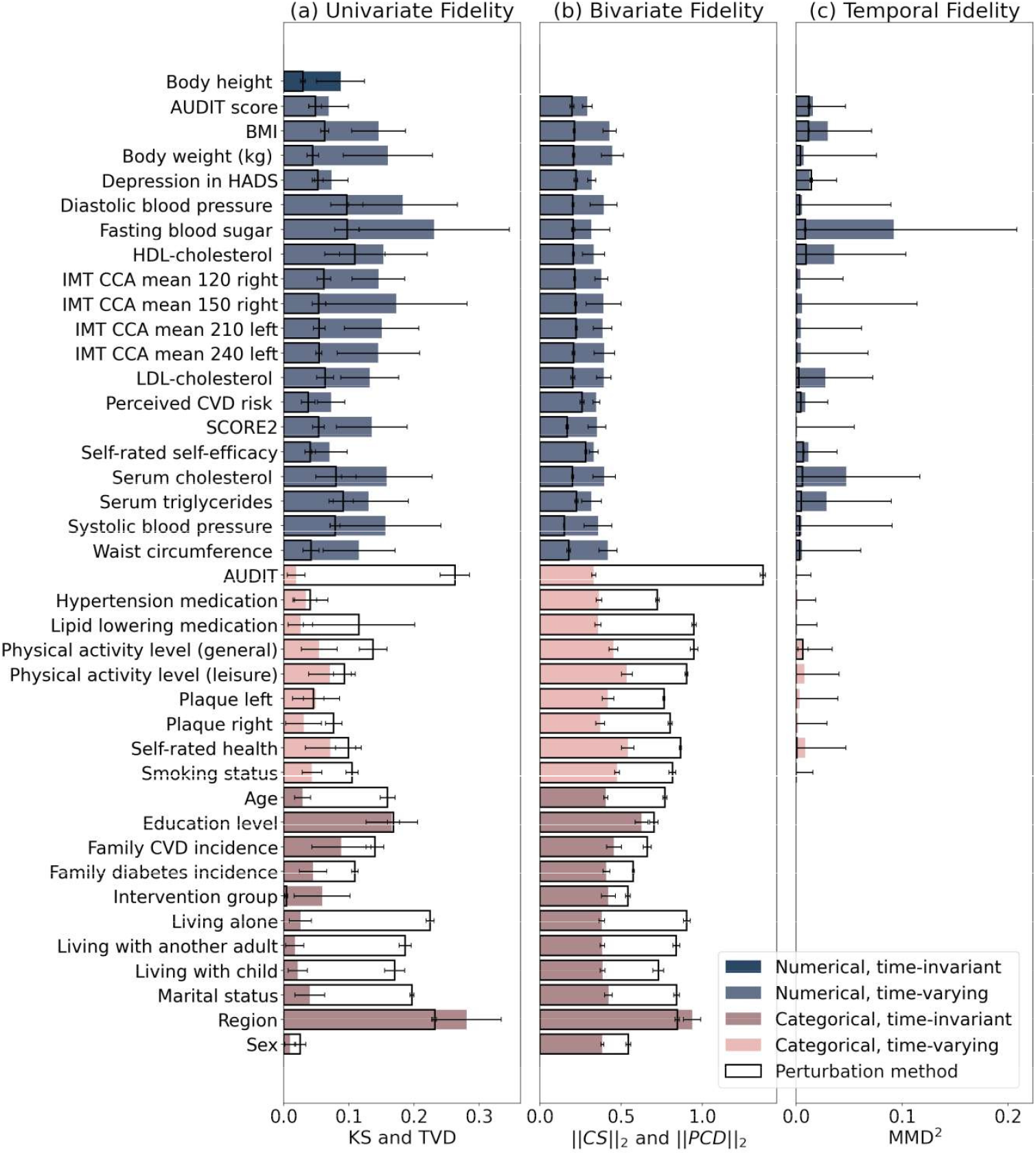
Variable-wise fidelity. Univariate, bivariate and temporal fidelity per variable of CTGAN and baseline. Univariate fidelity is reported in (a) and was computed using Kolmogorov-Smirnov statistic (*KS*) and total variation distance (*TV D*) for numerical and categorical variables respectively. Bivariate fidelity is reported in (b) and was computed using L2-norm of difference in Pearson correlation coefficient (||*PCD*||_2_) and L2-norm of contingency similarity complement (||*CS*^c^||_2_) for numerical and categorical variables respectively. Temporal fidelity is reported in (c) and was computed using mean maximum discrepancy squared (*MMD*^2^) for time-varying variables. Time-invariant variables are denoted with lighter colour shades for univariate and bivariate fidelity. Time-varying variables are reported by the mean across all time points. Lower values indicate better performance for all metrics. Note that *X*^ti^ only contains one numerical variable (*Height*) and was thus excluded from the plot.

For univariate numerical fidelity, CTGAN generally performed better for discrete numerical variables such as *AUDIT score, Depression in HADS* and *Self-rated self-efficacy* and the results for CTGAN are seen to be approaching the perturbation method results for these examples. For univariate categorical fidelity, CTGAN performed better than or equal to the perturbation method across all variables but *Region*, with the largest differences being observed for variables such as *AUDIT, Lipid lowering medication, Age, Marital status* and variables regarding living situation.

For bivariate fidelity, CTGAN exhibited stable performance across all variables, apart from *Region* and *Education level* for which performance was poorer. Note that the time-invariant variable *Body height (cm)* was excluded since it is the only numerical time-invariant variable.

The temporal fidelity of CTGAN showed more variance between variables than the perturbation method, which generally showed low values. Especially *Fasting blood sugar* followed by *Serum cholesterol* showed high *MMD*^2^ for CTGAN. This trend was not consistent across the univariate and bivariate fidelity.

### Stratified analysis

The fidelity results of the age-stratified analysis are shown in Fig 5. There was a small difference between the 60 year-olds and the complete dataset. Contrasting the 60 year-olds, there was an increase in distributional distances with decreasing age. The 40-year-olds showed a large increase in distributional distances in comparison to both 60-year-olds and 50-year-olds, and an overall high variance in fidelity despite the lower number of samples in the subset (7% of the original dataset).

**Fig 5.** Age-stratified fidelity. Fidelity for CTGAN stratified on age. The univariate fidelity includes mean Kolmogorov-Smirnov 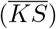 and mean total variation distance 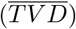 for numerical and categorical variables respectively. The bivariate fidelity includes mean L2-norm of difference in Pearson correlation coefficient 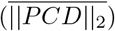 and mean L2-norm of contingency similarity complement 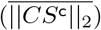 for numerical and categorical variables respectively. Note that *X*^ti^ only contains one numerical variable and was thus excluded from plot. Temporal fidelity is computed using mean maximum mean discrepancy squared 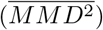 across all points in time. Lower values indicate better performance of all metrics.

The privacy results of the stratified analysis are reported in Table 2. A slightly higher protection of privacy was observed for the subsets, but the differences from the full dataset were, overall, relatively small.

**Table 2.** Age-stratified privacy.

| Privacy Metric | All | Age 40 | Age 50 | Age 60 |
| --- | --- | --- | --- | --- |
| $NNDR$ (↑) | 0.94 (0.00) | 0.94 (0.01) | 0.94 (0.00) | 0.94 (0.00) |
| $MDCR$ (↑) | 1.09 (0.012) | 1.15 (0.07) | 1.08 (0.02) | 1.10 (0.02) |
| ε-IR (↓) | 2.2% (0.3%) | 1.0% (0.7%) | 1.7% (1.5%) | 1.5% (0.4%) |

Privacy metrics of CTGAN with stratified results by age group.

## Discussion

This study primarily evaluated the fidelity and privacy of two GAN models, TGAN and CTGAN, compared to a rudimentary perturbation method on longitudinal data within long-term cardiovascular disease prevention. This was done to challenge the structural limitations of cross-sectional SDG methods and establish a benchmark for SDG methods on long-term longitudinal prevention data with inherent low covariance. The main findings were the following. CTGAN performed better than TGAN with consistent performance across fidelity metrics. Both models displayed higher privacy preservation than the perturbation method. CTGAN showed lower performance in numerical and temporal fidelity in reference to the perturbation method. CTGAN also showed lower performance within fidelity for younger age groups (e.g. 40-year-olds) than older age groups (e.g. 60-year-olds), as shown by the stratified analysis.

### Performance

Although CTGAN performed better than TGAN within fidelity, it varied against the perturbation method. CTGAN performed better than the perturbation method for categorical variables. The model had lower performance for numerical variables, specifically continuous numerical variables, in comparison to the perturbation method. The stronger categorical performance is consistent with previous results for CTGAN and DP-CGANS (also using a conditional vector) [21]. Furthermore, the results of CTGAN displayed large variation in the temporal domain. This may be due to large variable-wise variation in temporal complexity or stochastic variation from the sampling in the metric computation. The observed variation could not be conclusively attributed to a single factor and thus warrants further investigation.

The GAN models better preserved the privacy of the original samples than the perturbation method. This result differs from studies reporting that perturbation or obfuscation methods can outperform SDG methods, including CTGANs [30, 31], but can depend on the perturbation method of choice.

Differences in privacy between the GAN models and the perturbation method was observed for all metrics. *ε ™ IR* indicates the ratio of real samples that have a closer match in the synthetic dataset than the next real sample and should be below 9% [48]. TGAN and CTGAN performed within this threshold, whilst the perturbation method had a high percentage of 91.3%. Membership disclosure is not usually the most critical area of privacy since an attacker must already have a-priori information about the sample to identify its inclusion; however, identification may infer additional information or aid an attacker in recovering the encryption key if a weak encryption method is applied, thus risking the privacy of the entire cohort.

Identity disclosure is usually more harmful for individual participants since it deals with sample-wise distances, but can be a risk to the integrity of the entire cohort. *NNDR* has no general threshold levels, but a higher value means better protection. TGAN and CTGAN (0.94 for both) showed stronger results than the perturbation method (0.79), whose value was selected as the optimal point to avoid loss of fidelity. *MDCR* is upward unbounded, but a value below 1 indicates that samples in the real data more closely resemble the synthetic data than the other samples in the real data [48]. All models exceeded 1.

### Clinical aspects

This study highlighted clinical aspects in terms of (1) usability of the dataset and (2) availability of clinical datasets.

#### Usability of the synthetic data

Datasets within health and lifestyle contain multiple subgroups with different characteristics, often imbalanced. CTGAN is developed to handle this using a conditional sampling approach during training [29]. To evaluate its performance on imbalanced subgroups, a stratified analysis across age groups was performed. No notable differences in privacy were observed between the subgroups as all were within recommended thresholds; however, the same was not observed for fidelity. The fidelity results for the 60 year-olds, comprising 67% of the dataset, were similar to that of the entire dataset. Meanwhile, the results for 40- and 50-year-olds, comprising 7% and 26% respectively, showed greater distributional distances than the 60-year-olds for all fidelity metrics. This may be due to the imbalance or due to the different characteristics of the groups resulting from the different inclusion criteria in the VIPVIZA study.

The lower performance in younger individuals limits its application in an early prevention setting, which is an essential part of the clinical uniqueness of the VIPVIZA data. If the synthetic data set is used as a substitute for real data in the early phases of clinical research projects within prevention or early intervention, it may promote a premature and possibly untrue conclusion about the lack of signal for 40-year-olds. This must be acknowledged before the synthetic data is used and the primary intended use of the synthetic data should be focused on research within intervention towards people aged 50 and older.

The stratified analysis focused on the age groups, however, both age and sex are important variables for stratification to ensure equality across groups with known differences in health profiles. For this reason, a complementary stratified analysis was performed for sex and the results are reported in the supplementary materials S2. Within fidelity, CTGAN showed slightly lower performance for each subgroup compared to the results for the entire cohort. However, for both fidelity and privacy, there was no notable difference between the female and male subgroup. This may be because the sex variable was more balanced than the age variable (53% female, 47% male) or due to more pronounced differences in health between ages than between sex in this dataset.

Another common challenge with longitudinal health data concerns changes in the collection of the dataset. This can occur due to drift in study design over time, making the dataset more difficult to draw conclusions from [49]. In the VIPVIZA study, the variable *Depression in HADS* was introduced after the baseline measurement was initiated, explaining an initial large proportion of missing (33% for baseline). However, in this study, it did not affect the results since both CTGAN and the perturbation method showed similar performance on the variable as other variables. Other longitudinal datasets may be more prone to changes in study design over time, making the task more complex.

#### Implications on availability of data

Data sharing of clinical trial data is often regulated by controlled access, which refers to the sharing of protected data using contracts such as agreements or identification and vetoing of the data requester [19]. This limits the accessibility of the data, in favour of the integrity of participating individuals. The results of this study show that privacy can be preserved whilst maintaining fidelity when using CTGAN, which can reduce the need for controlled access when the risk is acceptable and thus increase the availability of clinical datasets. A semi-open access model with restraints in combination with synthetic data may be a first step towards making datasets more available for research, development and education.

### Limitations and future work

We have identified several limitations in this study that affect the generalisability of the results and methods to other datasets and applications. These include choices regarding the study design and the VIPVIZA dataset and methodological limitations.

#### Limitations of study design and the VIPVIZA dataset

In the VIPVIZA data, the proportion of missing data was generally low. This is not usually the case for conventional EHR, and the extendability of the results to other clinical data sets may therefore be affected. Furthermore, EHR are often naturally fragmented as different patients undergo different examinations. Fragmented datasets should be addressed with specialised imputation methods such as FragmGAN [50] and it should be investigated whether new methods such as attention based transformer prediction methods [51, 52] can be extended to a generative application and thus exclude the need for imputation entirely for fragmented datasets in SDG. For this reason, the results of this study on a structured dataset may not be generalisable to other longitudinal datasets within conventional EHR.

Another limitation concerns an assessment of utility since this study focused on the preservation of important characteristics of clinically relevant variables. For the VIPVIZA dataset, a more thorough evaluation of utility by assessment of clinical relevance of the synthetic data for a variety of tasks can involve reproducing previously performed clinical research with the synthetic dataset, then evaluate whether the findings are consistent with those obtained from the real data. In this work, the stratified analyses and the variable-wise fidelity serve as an indicator of whether the synthetic data could be used in place of the real data, before access to real data can be granted, but a more extensive evaluation should be explored in a separate study.

Additionally, we only included complete longitudinal cases in this study, which can introduce bias. The reasons for loss to follow up in VIPVIZA is reported in [42, 43], and a statistical analysis is reported in supplementary table 1 of [43], in which the mean SCORE2 is reportedly higher for participants with loss to follow up than longitudinal participants (5.34 (3.50) and 5.00 (3.10) respectively in the intervention group and 5.36 (3.49) and 5.08 (2.97) respectively in the control group). This shows that there are differences in risk level between the cohorts, and consequently, the covariate structures related to cardiovascular risk may be weaker in the longitudinal cohort than in the participants with loss to follow up.

#### Methodological limitations

We used a simple, yet common approach to sharing sensitive data by method of perturbation as reference. Since the method is inherently different from TGAN and CTGAN in the sense that it modifies variable values instead of having to learn the intrinsic covariate structures of the data, the comparison of performance in fidelity and privacy is limited. The perturbation method preserves values and longitudinal trajectories by default, and the performance in temporal fidelity is thus expected. Similarly, the privacy risk can be attributed to the multivariate characteristics being maintained despite perturbation, and this should be acknowledged when comparing the methods.

Another methodological limitation concerns the choice to not perform hyperparameter optimisation. Although we acknowledge that model tuning could lead to better results, the No Free Lunch Theorems for Optimisation tell us that all configurations perform equally well when averaged over all possible experiments [53]. As basis for comparison, we therefore preferred to maintain the baseline parameter set, so that the only way a method can outperform another one relies upon its specialisation to the structure of the problem under consideration [54]. Furthermore, in a framework where a model is not tuned, the winning method tends to correspond to the most robust, which is also a desirable characteristic [55].

Moreover, measurements that are equally spaced in time, as performed in the VIPVIZA study, are in general rare in conventional EHR. For example, hospital visits can be irregular [39], or the data from wearable devices is not usually consistent for personal or technical reasons [18, 56]. In such cases, the method of concatenation as employed in this study cannot be employed. Sequential methods for longitudinal SDG can better account for this irregularity [33, 35, 57], and future studies should explore a sequential approach for long-term prevention data. A sequential approach would also not require the exclusion of samples with loss to follow-up as the number of visits can vary.

## Conclusion

In this paper, we showed that CTGAN had better performance within fidelity compared to TGAN and a perturbation method on a long-term longitudinal dataset within cardiovascular disease prevention. Both TGAN and CTGAN exceeded the reference perturbation method in terms of privacy preservation, despite some previous studies reporting better performance for perturbation and obfuscation methods.

A more detailed assessment of CTGAN showed lower performance within fidelity on continuous numerical variables as well as temporal fidelity, in comparison to a reference perturbation method.

Finally, a stratified analysis on CTGAN showed variation in distributional distances across imbalanced age groups, highlighting the need for methods that ensure representation of known subgroups during training with techniques such as weighting. To the best of our knowledge, this is the first study evaluating CTGAN on a longitudinal dataset.

## Supporting information

S2 - Sex stratification

S1 - Descriptive statistics

## Supporting information

**S1 File. Descriptive statistics**. Descriptive statistics of variables in dataset.

**S2 File. Sex stratification**. Fidelity and privacy for CTGAN stratified on sex.

## Acknowledgments

Not applicable.

## References

1. World Heart Federation Geneva S. World Heart Report 2023: Confronting the World’s Number One Killer.; 2023.

2. Khot UN, Khot MB, Bajzer CT, Sapp SK, Ohman EM, Brener SJ, et al. Prevalence of conventional risk factors in patients with coronary heart disease. Jama. 2003;290(7):898–904.

3. Khera AV, Emdin CA, Drake I, Natarajan P, Bick AG, Cook NR, et al. Genetic risk, adherence to a healthy lifestyle, and coronary disease. New England Journal of Medicine. 2016;375(24):2349–58.

4. Webster R, Heeley E. Perceptions of risk: understanding cardiovascular disease. Risk management and healthcare policy. 2010:49–60.

5. Näslund U, Ng N, Lundgren A, Fhärm E, Grönlund C, Johansson H, et al. Visualization of asymptomatic atherosclerotic disease for optimum cardiovascular prevention (VIPVIZA): a pragmatic, open-label, randomised controlled trial. The Lancet. 2019;393(10167):133–42.

6. Blomstedt Y, Norberg M, Stenlund H, Nyström L, Lönnberg G, Boman K, et al. Impact of a combined community and primary care prevention strategy on all-cause and cardiovascular mortality: a cohort analysis based on 1 million person-years of follow-up in Västerbotten County, Sweden, during 1990–2006. BMJ open. 2015;5(12):e009651.

7. Asri H, Mousannif H, Al Moatassime H, Noel T. Big data in healthcare: Challenges and opportunities. In: 2015 International Conference on Cloud Technologies and Applications (CloudTech). IEEE; 2015. p. 1–7.

8. Pedroso AF, Khera R. Leveraging AI-enhanced digital health with consumer devices for scalable cardiovascular screening, prediction, and monitoring. npj Cardiovascular Health. 2025;2(1):34.

9. Martinez-Velazquez R, Gamez R, El Saddik A. Cardio Twin: A Digital Twin of the human heart running on the edge; 2019. p. 1–6. doi:10.1109/MeMeA.2019.8802162.

10. Mulder ST, Omidvari AH, Rueten-Budde AJ, Huang PH, Kim KH, Bais B, et al. Dynamic Digital Twin: Diagnosis, Treatment, Prediction, and Prevention of Disease During the Life Course. J Med Internet Res. 2022 Sep;24(9):e35675. Available from: https://www.jmir.org/2022/9/e35675. doi:10.2196/35675.

11. Akinosun AS, Polson R, Diaz-Skeete Y, De Kock JH, Carragher L, Leslie S, et al. Digital technology interventions for risk factor modification in patients with cardiovascular disease: systematic review and meta-analysis. JMIR mHealth and uHealth. 2021;9(3):e21061.

12. Gonzales A, Guruswamy G, Smith SR. Synthetic data in health care: A narrative review. PLOS Digital Health. 2023;2(1):e0000082.

13. Coorey G, Figtree GA, Fletcher DF, Snelson VJ, Vernon ST, Winlaw D, et al. The health digital twin to tackle cardiovascular disease—a review of an emerging interdisciplinary field. NPJ digital medicine. 2022;5(1):126.

14. El Emam K, Rodgers S, Malin B. Anonymising and sharing individual patient data. bmj. 2015;350.

15. El Emam K, Jonker E, Arbuckle L, Malin B. A systematic review of re-identification attacks on health data. PloS one. 2011;6(12):e28071.

16. Voigt P, Von dem Bussche A. The eu general data protection regulation (gdpr). A practical guide, 1st ed, Cham: Springer International Publishing. 2017;10(3152676):10–5555.

17. for Disease Control c, Prevention, et al. HIPAA privacy rule and public health. Guidance from CDC and the US Department of Health and Human Services. MMWR: Morbidity and mortality weekly report. 2003;52(Suppl 1):1–17.

18. Brown R, Coventry L, Sillence E, Blythe J, Stumpf S, Bird J, et al. Collecting and sharing self-generated health and lifestyle data: Understanding barriers for people living with long-term health conditions–a survey study. Digital health. 2022;8:20552076221084458.

19. Rodriguez A, Tuck C, Dozier MF, Lewis SC, Eldridge S, Jackson T, et al. Current recommendations/practices for anonymising data from clinical trials in order to make it available for sharing: A scoping review. Clinical Trials. 2022;19(4):452–63.

20. Choi E, Biswal S, Malin B, Duke J, Stewart WF, Sun J. Generating multi-label discrete patient records using generative adversarial networks. In: Machine learning for healthcare conference. PMLR; 2017. p. 286–305.

21. Sun C, van Soest J, Dumontier M. Generating synthetic personal health data using conditional generative adversarial networks combining with differential privacy. Journal of Biomedical Informatics. 2023;143:104404.

22. Little RJ. Statistical analysis of masked data. Journal of Official statistics. 1993;9(2):407.

23. Goodfellow IJ, Pouget-Abadie J, Mirza M, Xu B, Warde-Farley D, Ozair S, et al. Generative adversarial nets. In: Proceedings of the 28th International Conference on Neural Information Processing Systems - Volume 2. NIPS’14. Cambridge, MA, USA: MIT Press; 2014. p. 2672–2680.

24. Mirza M, Osindero S. Conditional Generative Adversarial Nets. CoRR. 2014;abs/1411.1784. Available from: http://arxiv.org/abs/1411.1784. 1411.1784.

25. Arjovsky M, Bottou L. Towards Principled Methods for Training Generative Adversarial Networks; 2017. Available from: https://arxiv.org/abs/1701.04862. 1701.04862.

26. Arjovsky M, Chintala S, Bottou L. Wasserstein Generative Adversarial Networks. In: Precup D, Teh YW, editors. Proceedings of the 34th International Conference on Machine Learning. vol. 70 of Proceedings of Machine Learning Research. PMLR; 2017. p. 214–23. Available from: https://proceedings.mlr.press/v70/arjovsky17a.html.

27. Gulrajani I, Ahmed F, Arjovsky M, Dumoulin V, Courville A. Improved Training of Wasserstein GANs; 2017. Available from: https://arxiv.org/abs/1704.00028. 1704.00028.

28. Xu L, Veeramachaneni K. Synthesizing Tabular Data using Generative Adversarial Networks. arXiv e-prints. 2018 Nov:1811.11264. 1811.11264. doi:10.48550/arXiv.1811.11264.

29. Xu L, Skoularidou M, Cuesta-Infante A, Veeramachaneni K. Modeling tabular data using conditional gan. Advances in neural information processing systems. 2019;32.

30. Marino S, Cassidy R, Nanni J, Wang Y, Liu Y, Tang M, et al. Medical data sharing and synthetic clinical data generation–maximizing biomedical resource utilization and minimizing participant re-identification risks. NPJ Digital Medicine. 2025;8(1):526.

31. Francis P, Jurak G, Leskošek B, Otte K, Prasser F. Comparison of three anonymization tools for a Health Fitness Study. Scientific Data. 2025;12(1):1548.

32. Ahmed HA, Nepomuceno JA, Vega-Márquez B, Nepomuceno-Chamorro IA. Synthetic Data Generation for Healthcare: Exploring Generative Adversarial Networks Variants for Medical Tabular Data. International Journal of Data Science and Analytics. 2025:1–16.

33. Zhang Z, Yan C, Lasko TA, Sun J, Malin BA. SynTEG: a framework for temporal structured electronic health data simulation. Journal of the American Medical Informatics Association. 2021;28(3):596–604.

34. Li J, Cairns BJ, Li J, Zhu T. Generating synthetic mixed-type longitudinal electronic health records for artificial intelligent applications. NPJ digital medicine. 2023;6(1):98.

35. Theodorou B, Xiao C, Sun J. Synthesize high-dimensional longitudinal electronic health records via hierarchical autoregressive language model. Nature communications. 2023;14(1):5305.

36. Wang Y, Xu X, Jin T, Li X, Xie G, Wang J. Inpatient2vec: Medical representation learning for inpatients. In: 2019 IEEE International Conference on Bioinformatics and Biomedicine (BIBM). IEEE; 2019. p. 1113–7.

37. Wang JL, Chiou JM, Müller HG. Functional data analysis. Annual Review of Statistics and its application. 2016;3(1):257–95.

38. Philipps V, Freedman L, Deffner V, Helmer C, Boshuizen H, Thiébaut A, et al. Including an infrequently measured time-dependent error-prone covariate in survival analyses: a simulation-based comparison of methods. arXiv preprint 250216362. 2025.

39. Perkonoja K, Auranen K, Virta J. Methods for generating and evaluating synthetic longitudinal patient data: a systematic review. arXiv preprint 230912380. 2023.

40. Bengtsson A, Norberg M, Ng N, Carlberg B, Grönlund C, Hultdin J, et al. The beneficial effect over 3 years by pictorial information to patients and their physician about subclinical atherosclerosis and cardiovascular risk: Results from the VIPVIZA randomized clinical trial. American Journal of Preventive Cardiology. 2021;7:100199. Available from: https://www.sciencedirect.com/science/article/pii/S2666667721000532. doi:10.1016/j.ajpc.2021.100199.

41. Norberg M, Wall S, Boman K, Weinehall L. The Västerbotten Intervention Programme: background, design and implications. Global health action. 2010;3(1):4643.

42. Bengtsson A, Norberg M, Ng N, Carlberg B, Grönlund C, Hultdin J, et al. The beneficial effect over 3 years by pictorial information to patients and their physician about subclinical atherosclerosis and cardiovascular risk: results from the VIPVIZA randomized clinical trial. American journal of preventive cardiology. 2021;7:100199.

43. Norberg M, Wennberg P, Wester P, Själander A, Liv P. Evaluation after delayed and repeated intervention in the VIPVIZA-extended randomized controlled trial: beneficial results 6 years after baseline. European Heart Journal Open. 2026;6(2):oeag047.

44. Hernandez M, Epelde G, Alberdi A, Cilla R, Rankin D. Synthetic data generation for tabular health records: A systematic review. Neurocomputing. 2022;493:28–45.

45. Kaabachi B, Despraz J, Meurers T, Otte K, Halilovic M, Kulynych B, et al. A scoping review of privacy and utility metrics in medical synthetic data. NPJ digital medicine. 2025;8(1):60.

46. Ghosheh GO, Li J, Zhu T. A survey of generative adversarial networks for synthesizing structured electronic health records. ACM Computing Surveys. 2024;56(6):1–34.

47. Synthetic Data Metrics; 2023. Version 0.12.0. Available from: https://docs.sdv.dev/sdmetrics/.

48. Lautrup AD, Hyrup T, Zimek A, Schneider-Kamp P. Syntheval: a framework for detailed utility and privacy evaluation of tabular synthetic data. Data Mining and Knowledge Discovery. 2024;39(1). doi:10.1007/s10618-024-01081-4.

49. Matton L, Beunen G, Duvigneaud N, Wijndaele K, Philippaerts R, Claessens A, et al. Methodological issues associated with longitudinal research: findings from the Leuven Longitudinal Study on Lifestyle, Fitness and Health (1969–2004). Journal of sports sciences. 2007;25(9):1011–24.

50. Fang F, Bao S. FragmGAN: generative adversarial nets for fragmentary data imputation and prediction. Statistical Theory and Related Fields. 2024;8(1):15–28.

51. Caruso CM, Soda P, Guarrasi V. Not Another Imputation Method: A Transformer-based Model for Missing Values in Tabular Datasets. arXiv preprint 240711540. 2024.

52. Caruso CM, Soda P, Guarrasi V. MARIA: a multimodal transformer model for incomplete healthcare data. Computers in Biology and Medicine. 2025;196:110843.

53. Wolpert DH, Macready WG. No free lunch theorems for optimization. IEEE transactions on evolutionary computation. 2002;1(1):67–82.

54. Ho YC, Pepyne DL. Simple explanation of the no-free-lunch theorem and its implications. Journal of optimization theory and applications. 2002;115(3):549–70.

55. Fernández A, López V, Galar M, Del Jesus MJ, Herrera F. Analysing the classification of imbalanced data-sets with multiple classes: Binarization techniques and ad-hoc approaches. Knowledge-based systems. 2013;42:97–110.

56. Cho S, Ensari I, Weng C, Kahn MG, Natarajan K. Factors affecting the quality of person-generated wearable device data and associated challenges: rapid systematic review. JMIR mHealth and uHealth. 2021;9(3):e20738.

57. Yoon J, Jarrett D, Van der Schaar M. Time-series generative adversarial networks. Advances in neural information processing systems. 2019;32.

