## Supplementary material for "Evaluation of synthetic data generation for longitudinal 6-year follow-up subclinical cardiovascular disease data using GAN models": S2 - Sex stratification

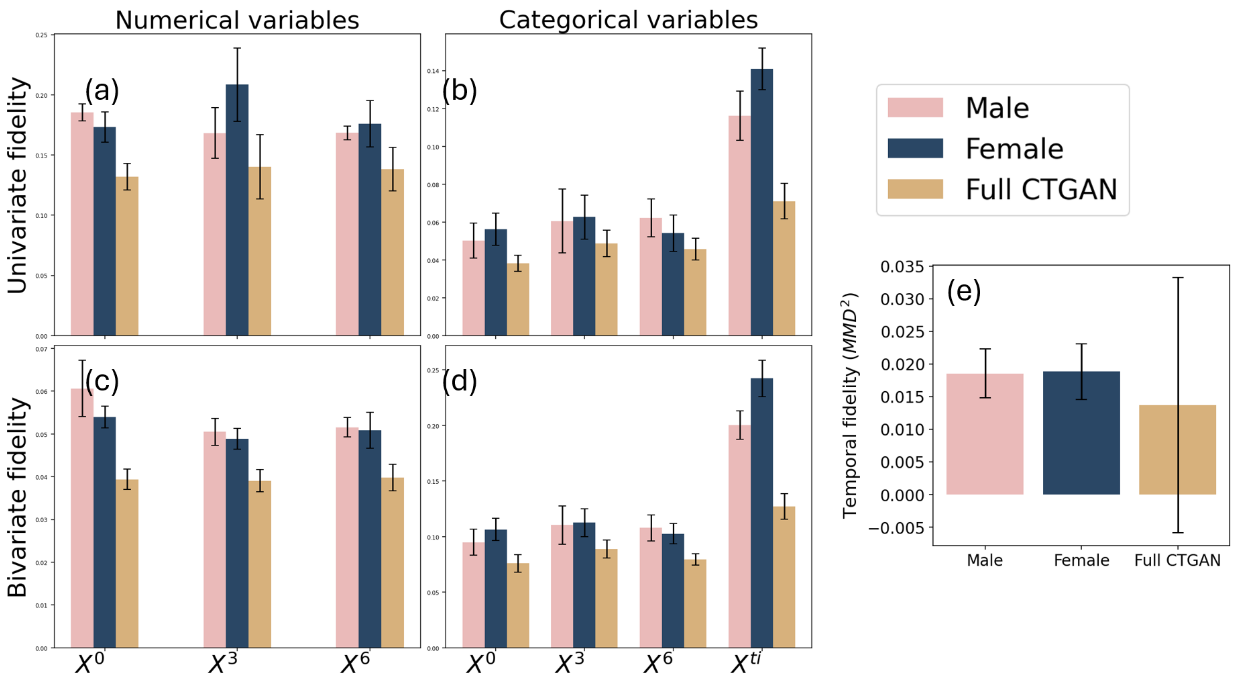


Figure 1: Fidelity for CTGAN stratified on sex. The univariate fidelity includes mean Kolmogorov-Smirnov ($\overline{KS}$) and mean total variation distance ($\overline{TVD}$) for numerical and categorical variables respectively. The bivariate fidelity includes mean L2-norm of difference in Pearson correlation coefficient ($\overline{||PCD||_2}$) and mean L2-norm of contingency similarity complement ($\overline{||CS^c||_2}$) for numerical and categorical variables respectively. Note that $X^{ti}$ only contains one numerical variable and was thus excluded from plot. Temporal fidelity is computed using mean maximum mean discrepancy squared ($\overline{MMD^2}$) across all points in time. Lower values indicate better performance of all metrics.

Table 1: Privacy metrics of CTGAN with stratified results by sex.

| **Privacy Metric** | **All** | **Male** | **Female** |
|---|---|---|---|
| $NNDR$ (↑) | 0.94 (0.00) | 0.94 (0.00) | 0.94 (0.00) |
| $MDCR$ (↑) | 1.09 (0.02) | 1.09 (0.03) | 1.09 (0.01) |
| $\varepsilon$-IR (↓) | 2.1% (0.3%) | 1.3% (0.3%) | 1.4% (0.3%) |
