## Supplementary material for "Evaluation of synthetic data generation for longitudinal 6-year follow-up subclinical cardiovascular disease data using GAN models": S1 - Descriptive statistics

### S1 Descriptive statistics

#### S1.1 Time-invariant variables

| Variable | Level | Observed | Imputed | Missing |
|---|---|---|---|---|
| Age | 60 years | 66.6% | – | – |
| | 50 years | 26.5% | – | |
| | 40 years | 6.9% | – | |
| Marital status | Partner | 79.8% | 79.9% | 1.03% |
| | Single | 10.2% | 10.2% | |
| | Other | 10.0% | 10.0% | |
| Presently living with: Adult | False | 17.9% | – | – |
| | True | 82.1% | – | |
| Presently living with: Child | False | 71.5% | – | – |
| | True | 28.5% | – | |
| Presently living with: I live alone | False | 87.1% | – | – |
| | True | 12.9% | – | |
| Family CVD incidence | No | 68.6% | 68.5% | 3.56% |
| | Yes | 31.4% | 31.5% | |
| Family diabetes incidence | No | 70.6% | 70.7% | 2.83% |
| | Yes | 29.4% | 29.3% | |
| Education level | Basic | 7.7% | 7.7% | 0.84% |
| | Secondary | 55.8% | 55.8% | |
| | Higher | 36.5% | 36.5% | |
| Intervention group | False | 49.5% | – | – |
| | True | 50.5% | – | |
| Body height (cm)* | – | 171.65 (9.52) | 171.65 (9.52) | 0.04% |
| Region | Umeå | 41.1% | – | – |
| | Rural inland | 21.5% | – | |
| | Rural coastland | 20.4% | – | |
| | Skellefteå | 17.0% | – | |
| Sex | False | 53.0% | – | – |
| | True | 47.0% | – | |

Table 1: Descriptive statistics of time-invariant variables. Label proportions are reported for categorical variables and mean (SD) is reported for numerical variables (indicated by ”*”). The statistics are reported for observed and imputed variables, as well as proportion of missing data. For the imputed data, the mean statistic across all versions of imputation is reported.

#### S1.2 Time-varying variables

##### S1.2.1 Numerical variables

| | Baseline | | | Year 3 | | | Year 6 | | |
|---|---|---|---|---|---|---|---|---|---|
| **Variable** | **Observed** | **Imputed** | **Missing** | **Observed** | **Imputed** | **Missing** | **Observed** | **Imputed** | **Missing** |
| AUDIT score | 3.22 (2.49) | 3.22 (2.48) | 1.99% | 3.15 (2.52) | 3.15 (2.52) | **7.01%** | 3.09 (2.57) | 3.09 (2.57) | **10.18%** |
| Fasting blood sugar (mmol/l) | 5.35 (1.16) | 5.35 (1.16) | 0.27% | 5.50 (1.06) | 5.50 (1.06) | 1.72% | 5.53 (1.03) | 5.53 (1.02) | 1.88% |
| BMI | 27.34 (4.63) | 27.35 (4.63) | 0.08% | 27.63 (4.80) | 27.62 (4.80) | 1.49% | 27.72 (4.93) | 27.73 (4.93) | 1.19% |
| Diastolic blood pressure (mmHg) | 82.17 (10.16) | 82.18 (10.16) | 0.08% | 85.52 (9.81) | 85.52 (9.82) | 1.80% | 85.24 (9.50) | 85.25 (9.52) | 1.11% |
| Depression in HADS | 2.71 (2.45) | 2.70 (2.44) | **32.96%** | 2.64 (2.67) | 2.65 (2.67) | 2.41% | 2.62 (2.61) | 2.61 (2.61) | 4.63% |
| HDL-cholesterol (mmol/l) | 1.40 (0.42) | – | – | 1.55 (0.45) | 1.55 (0.45) | 1.49% | 1.56 (0.44) | 1.56 (0.44) | 1.00% |
| IMT CCA mean 210 left | 0.66 (0.14) | 0.66 (0.14) | 0.77% | 0.67 (0.15) | 0.67 (0.15) | 0.31% | 0.68 (0.16) | 0.68 (0.16) | 0.54% |
| IMT CCA mean 240 left | 0.67 (0.15) | 0.67 (0.15) | 0.46% | 0.68 (0.16) | 0.68 (0.16) | 0.31% | 0.69 (0.16) | 0.69 (0.16) | 0.57% |
| IMT CCA mean 120 right | 0.66 (0.14) | 0.66 (0.14) | 0.61% | 0.67 (0.15) | 0.67 (0.15) | 0.34% | 0.69 (0.16) | 0.69 (0.16) | 0.44% |
| IMT CCA mean 150 right | 0.65 (0.13) | 0.65 (0.13) | 0.69% | 0.66 (0.14) | 0.66 (0.14) | 0.38% | 0.67 (0.14) | 0.67 (0.14) | 0.46% |
| LDL-cholestero (mmol/l) | 3.57 (0.97) | 3.57 (0.97) | 2.07% | 3.23 (1.01) | 3.23 (1.01) | 2.49% | 2.84 (1.08) | 2.84 (1.08) | 2.26% |
| Waist circum-ference (cm) | 95.66 (12.71) | 95.65 (12.70) | 1.34% | 97.48 (12.96) | 97.47 (12.95) | 2.41% | 97.78 (13.47) | 97.77 (13.46) | 2.34% |
| Systolic blood pressure (mmHg) | 128.66 (15.81) | – | – | 129.84 (16.05) | 129.85 (16.07) | 1.80% | 134.39 (16.40) | 134.38 (16.40) | 1.11% |
| SCORE2 | 0.05 (0.03) | 0.05 (0.03) | 0.69% | 0.06 (0.03) | 0.06 (0.03) | 2.18% | 0.07 (0.04) | 0.07 (0.04) | 2.14% |
| Serum cholesterol (mmol/l) | 5.61 (1.08) | – | – | 5.39 (1.12) | 5.39 (1.12) | 1.49% | 5.02 (1.20) | 5.02 (1.20) | 1.00% |
| Perceived CVD risk | 4.03 (2.26) | 4.02 (2.26) | 1.91% | 4.72 (2.22) | 4.73 (2.22) | **10.53%** | 5.06 (2.16) | 5.06 (2.16) | **5.51%** |
| Self-rated self-efficacy | 7.42 (1.95) | 7.42 (1.94) | 1.88% | 7.12 (2.03) | 7.11 (2.04) | **10.49%** | 7.13 (2.12) | 7.13 (2.12) | **5.55%** |
| Serum triglycerides (mmol/l) | 1.46 (0.96) | – | – | 1.35 (0.74) | 1.35 (0.74) | 1.49% | 1.39 (0.82) | 1.39 (0.82) | 1.00% |
| Body weight (kg) | 80.79 (16.18) | 80.79 (16.18) | 0.08% | 81.50 (16.51) | 81.50 (16.50) | 1.49% | 81.54 (16.96) | 81.55 (16.97) | 1.19% |

Table 2: Descriptive statistics of time-varying numerical variables at each time point. Mean (SD) is reported for observed and imputed variables, along with proportion of missing data. Missing proportions greater than 5% are highlighted in bold.

###### S1.2.2 Categorical variables

| | | Baseline | | | Year 3 | | | Year 6 | | |
|---|---|---|---|---|---|---|---|---|---|---|
| **Variable** | **Level** | **Observed** | **Imputed** | **Missing** | **Observed** | **Imputed** | **Missing** | **Observed** | **Imputed** | **Missing** |
| AUDIT | Not at risk | 92.3% | 92.3% | 1.99% | 92.4% | 92.4% | 7.01% | 92.4% | 92.3% | 10.18% |
| | Risky alcohol use | 7.7% | 7.7% | | 7.6% | 7.6% | | 7.6% | 7.7% | |
| Hypertension medication | False | 69.7% | 69.7% | 3.29% | 60.4% | 60.4% | 0.69% | 52.6% | 52.6% | 0.84% |
| | True | 30.3% | 30.3% | | 39.6% | 39.6% | | 47.4% | 47.4% | |
| Lipid lowering medication | False | 88.3% | 88.2% | 3.29% | 77.2% | 77.2% | 0.69% | 57.3% | 57.3% | 0.88% |
| | True | 11.7% | 11.8% | | 22.8% | 22.8% | | 42.7% | 42.7% | |
| Self-rated health | Pretty good | 48.1% | 48.1% | 0.38% | 51.5% | 51.7% | 4.10% | 52.1% | 52.0% | **7.89%** |
| | Very good | 25.2% | 25.2% | | 19.0% | 18.9% | | 17.5% | 17.5% | |
| | Somewhat good | 19.9% | 19.9% | | 22.9% | 22.8% | | 23.2% | 23.3% | |
| | Bad / pretty bad | 6.8% | 6.8% | | 6.6% | 6.6% | | 7.2% | 7.2% | |
| Physical activity (leisure) | Light | 51.4% | 51.4% | 0.69% | 51.1% | 51.2% | 3.60% | 51.9% | 51.9% | **7.43%** |
| | Moderate | 27.4% | 27.4% | | 25.2% | 25.2% | | 23.2% | 23.3% | |
| | Regular exercise | 13.0% | 13.0% | | 16.1% | 16.0% | | 17.5% | 17.4% | |
| | Sedentary | 8.2% | 8.2% | | 7.5% | 7.5% | | 7.4% | 7.4% | |
| Physical activity (general) | Meets recommendations | 58.3% | 58.3% | 0.92% | 64.1% | 64.0% | 4.10% | 68.0% | 67.9% | **7.85%** |
| | Moderately active | 25.4% | 25.3% | | 21.0% | 21.0% | | 18.4% | 18.4% | |
| | Low activity | 16.3% | 16.4% | | 14.9% | 14.9% | | 13.6% | 13.6% | |
| Plaque left | False | 66.5% | – | – | 58.5% | 58.5% | 0.27% | 53.8% | 53.7% | 0.46% |
| | True | 33.5% | – | | 41.5% | 41.5% | | 46.2% | 46.3% | |
| Plaque right | False | 67.0% | 66.9% | 0.04% | 59.6% | 59.6% | 0.31% | 56.0% | 55.9% | 0.46% |
| | True | 33.0% | 33.0% | | 40.4% | 40.4% | | 44.0% | 44.1% | |
| Smoking | Never | 52.8% | 52.8% | 0.19% | 51.2% | 51.2% | 0.73% | 50.7% | 50.7% | 0.38% |
| | Former daily | 23.4% | 23.4% | | 24.4% | 24.4% | | 23.7% | 23.8% | |
| | Former occasional | 13.0% | 13.0% | | 15.3% | 15.4% | | 17.7% | 17.7% | |
| | Current | 10.9% | 10.8% | | 9.1% | 9.0% | | 7.9% | 7.9% | |

Table 3: Descriptive statistics of time-varying categorical variables at each time point. Label proportions are reported for observed and imputed variables, as well as proportion of missing data. The proportion of missing data is in bold for variables with ¿5% of missing. For the imputed data, the mean statistic across all versions of imputation is reported.
